# Microvascular Injury in Mild Traumatic Brain Injury Accelerates Alzheimer-like Pathogenesis in Mice

**DOI:** 10.1101/2020.04.12.036392

**Authors:** Yingxi Wu, Jianxiong Zeng, Brock Pluimer, Shirley Dong, Xiaochun Xie, Xinying Guo, Xinyan Liang, Sudi Feng, Haijian Wu, Youzhen Yan, Jian-Fu Chen, Naomi Sta Maria, Qingyi Ma, Fernando Gomez-Pinilla, Zhen Zhao

**Affiliations:** Center for Neurodegeneration and Regeneration, Zilkha Neurogenetic Institute and Department of Physiology and Biophysics, Keck School of Medicine, University of Southern California, Los Angeles, California, 90033, USA; Neuroscience Graduate Program, Keck School of Medicine, University of Southern California, Los Angeles, California, 90033, USA; Center for Craniofacial Molecular Biology, Herman Ostrow School of Dentistry, University of Southern California, Los Angeles, California, 90033, USA; Lawrence D. Longo, MD Center for Perinatal Biology, Division of Pharmacology, Department of Basic Sciences, Loma Linda University School of Medicine, Loma Linda, California, 92350, USA; Brain Injury Research Center and Department of Neurosurgery, University of California, Los Angeles, Los Angeles, California, 90095, USA

**Keywords:** Traumatic brain injury, Alzheimer’s disease, microvascular injury, blood-brain barrier, β-amyloid

## Abstract

**Introduction:** Traumatic brain injury (TBI) is considered as the most robust environmental risk factor for Alzheimer’s disease (AD). Besides direct neuronal injury and neuroinflammation, vascular impairment is also a hallmark event of the pathological cascade after TBI. However, the vascular connection between TBI and subsequent AD pathogenesis remains underexplored.

**Methods:** We established a closed-head mild TBI (mTBI) model in mice with controlled cortical impact, and examined the time courses of microvascular injury, blood-brain barrier (BBB) dysfunction, gliosis and motor function impairment in wild type C57BL/6 mice. We also determined the brain clearance of β-amyloid, as well as amyloid pathology and cognitive functions after mTBI in the 5xFAD mouse model of AD.

**Results:** mTBI induced microvascular injury with BBB breakdown, pericyte loss and cerebral blood flow reduction in mice, which preceded gliosis. mTBI also impaired brain amyloid clearance via the vascular pathways. More importantly, mTBI accelerated amyloid pathology and cognitive impairment in the 5xFAD mice.

**Discussion:** Our data demonstrated that microvascular injury plays a key role in the pathogenesis of AD after mTBI. Therefore, restoring vascular functions might be beneficial for patients with mTBI, and potentially reduce the risk of developing AD.

## Background

Alzheimer’s disease (AD) is an age-related progressive neurodegenerative condition, manifesting amyloid plaque and neurofibrillary tangle formation, neurovascular and neuroimmune dysfunctions, and cognitive impairment [1,2]. While advanced aging increases the likelihood of AD, genetic inheritance and environmental risk factors also contribute significantly [3]. For example, Traumatic brain injury (TBI) is considered as the most robust environmental risk factor for AD [4]. TBI is a leading cause of death and disability, particularly in young adults, resulting in a great impact on productivity and dependence on health care in later life [5,6]. Both clinical and preclinical studies have demonstrated that TBI triggers multiple neurodegenerative cascades, including axonal and dendritic damage, excitatory toxicity, neuroinflammation and cell death [7,8], as well as cerebrovascular impairment such as edema, circulatory insufficiency, and blood-brain barrier (BBB) breakdown [9,10], exhibiting a high similarity with AD [6,11]. TBI is highly prevalent during military service and contact sports, and doubles the risk of developing AD and dementia [12,13]. More importantly, it also exacerbates certain pathological events that are specific to AD, including the brain’s overproduction and accumulation of β-amyloid (Aβ), and neurofibrillary tangles consisting of hyperphosphorylated Tau [14]. Yet the underlying mechanisms remain elusive.

Histological and neuroimaging assessments have demonstrated that microvascular injury with BBB breakdown are common in TBI patients. It occurs during the acute/subacute phase of TBI [15,16] and may last for years in nearly 50% of the survivors [17,18], resulting in long-term brain dysfunctions. Such microvascular endophenotype was well recapitulated in animal models of TBI [6], with completely different injury settings including fluid percussion injury, controlled cortical impact and blast injury. Mechanistically, TBI induces endothelial dysfunction [19], disrupts the crosstalk between endothelial cells and pericytes [20], reduces cerebral blood flow (CBF) and causes tissue hypoxia [21], upregulation of vascular endothelial growth factor (VEGF) and metallopeptidases [22,23], leukocyte infiltration [24], gliosis and neuroinflammation [7], which all contribute to BBB dysfunctions.

As the interface between the circulation and central nervous system (CNS), the BBB plays a key role in normal brain physiological regulation and homeostasis [25]. Anatomically, it consists of continuous endothelia sealed by highly specialized intercellular junctional structures to restrict paracellular flow, and highly selective transport systems eliminating toxic substances yet allowing exchanges of nutrients and metabolites between circulation and brain [11,25]. Therefore, microvascular injury often results in a cascade of events including extravasation of plasma proteins that are potentially toxic to neuronal cells, parenchymal edema and hypoxia, metabolic stress including endoplasmic reticulum (ER) and mitochondrial stress, accumulation of metabolic wastes, activation of microglia and astrocytes, and eventually neuronal dysfunctions [11,25]. Although microvascular injury is a shared endophenotype between TBI and AD [6], and a strong modifier of AD pathogenesis and progression [11], the vascular link between TBI and AD has not been investigated in depth.

Since mild TBI (mTBI) represents nearly 85% of cases, we established a close-head mTBI model in rodents to investigate vascular impairment after mTBI, as well as to understand its link to AD pathogenesis. We found that mTBI induced microvascular injury and BBB dysfunction in the acute/subacute phase of mTBI, which was tightly associated with circulatory insufficiency and pericyte loss, and in general preceded astrogliosis and microglia activation. In addition, applying mTBI in 5xFAD transgenic model of AD demonstrated that mTBI accelerated microvascular injury, amyloid pathology and cognitive impairments, which may be attributed to the impairment of brain clearance of amyloid via the cerebrovascular system. Therefore, improving vascular functions and BBB integrity in patients with mTBI may be beneficial and even reduce the risk of developing AD.

## Methods

### Animals

12-16 weeks old C57BL/6 mice and 5xFAD mice were used. 5xFAD mouse model is a well-characterized mouse model of AD, with expression of human *APP* and *PSEN1* transgenes with a total of five AD-linked mutations: the Swedish (K670N/M671L), Florida (I716V), and London (V717I) mutations in APP, and the M146L and L286V mutations in PSEN1 [26]. All procedures are approved by the Institutional Animal Care and Use Committee at the University of Southern California using the US National Institutes of Health (NIH) guidelines. Since the previous report showed no sex difference in selected motor and cognitive impairment after mTBI between male and female mice [27], both male and female mice were used in this study.

### Randomization and Blinding

All animals were randomized for their genotype information and surgical procedures, and included in the analysis. The operators responsible for the experimental procedures and data analysis were blinded and unaware of group allocation throughout the experiments.

### Closed-head mTBI mouse model

Mice received a mild closed-head impact following the procedures previously reported with minor modifications [28]. Briefly, for a precise impact mimicking mTBI, we utilized a stereotactic frame with adjustable angle (Stereotaxic Alignment System, KOPF) to ensure the impacted surface (the impact central point at Bregma -2 mm and lateral 2.5 mm) of the skull was presented perpendicular to the impactor (**Sup Fig. 1B**, Brain and Spinal Cord Impactor, #68099, RWD Life Science). We also custom-made plastic flat-tip impactor (RWD Life Science) to reduce the impact. Mice were anesthetized with ketamine/xylazine (90 mg/kg, 9 mg/kg; i.p.), then the head was fixed with a 15° angle, and a gel pad was placed under the head in order to maximally avoid the skull fractures and intracerebral bleeding. An incision was made to expose skull surface after anesthesia and then the mice were subjected to an impact using a 4 mm plastic flat-tip impactor. The velocity was 3 m/s, the depth was 1.0 mm and the impact duration was 180 ms (see **Sup Fig.1C-D**). After the procedure, the mice were put back to their cages with heating to recover from the anesthesia. For sham-operated mice, the same procedure was performed as the mTBI group except the tip was set 2.0 mm above the skull.

### Open-head severe TBI (sTBI) mouse model

Mice received the sTBI following the procedures previously reported [29]. Briefly, mice were anesthetized with ketamine/xylazine (90 mg/kg, 9 mg/kg; i.p.), and the mouse head was then fixed in the stereotactic frame (Stereotaxic Alignment System, KOPF). A craniotomy was made using a drill and 4.5-mm trephine in the center between Lambda and Bregma, and then the bone flap was removed. Mice were subjected to an impact using a 4 mm metal flat-tip impactor (Brain and Spinal Cord Impactor, #68099, RWD Life Science). The impact central point was Bregma -2.5 mm and lateral 2.5 mm. The velocity was 3 m/s, the depth was 1 mm and the impact duration was 180 ms. Then the scalp was closed with suture, and the mice were put back to their cages to recover from the anesthesia. For sham-operated mice, the same procedure was performed as the sTBI group except the tip was set 2.0 mm above the skull.

## Behavioral Tests

### Rotarod Test

Mice were trained on an accelerating (5-20 rpm) rotating rod (rotarod) for 3 days before TBI. Test sessions consist of six trials at a variable speed (an initial velocity of 5 rpm was used for the first 10 s, a linear increase from 5 to 10 rpm for the next 30 s, and a linear increase from 10 to 20 rpm between 40 to 80 s). The final score was determined as the mean time that a mouse was able to remain on the rod over six trials.

### Foot Fault test

The foot-fault test was performed as previously described [30]. Mice were placed on hexagonal grids of different sizes. Mice placed their paws on the wire while moving along the grid. With each weight-bearing step, the paw may fall or slip between the wire. This was recorded as a foot fault. The total number of steps (movement of each forelimb) that the mouse used to cross the grid was counted, and the total number of foot faults for each forelimb was recorded.

### Contextual fear conditioning

A hippocampus-dependent fear conditioning test was performed as previously described [31]. The experiments were performed using standard conditioning chambers housed in a soundproof isolation cubicle and equipped with a stainless-steel grid floor connected to a solid-state shock scrambler. The scrambler was connected to an electronic constant-current shock source that was controlled via Freezeframe software (Coulbourne Instruments). A digital camera was mounted on the steel ceiling and behavior was monitored. During training, mice were placed in the conditioning chamber for 4 min and received two-foot shocks (0.25 mA, 2 s) at 1-min interval starting 2 min after placing the mouse in the chamber. Contextual memory was tested the next day in the chamber without foot shocks. Hippocampus-dependent fear memory formation was evaluated by scoring freezing behavior (the absence of all movement except for respiration). For the contextual fear conditioning paradigms, the automated Freezeframe system was used to score the percentage of total freezing time with a threshold set at 10% and minimal bout duration of 0.25 s.

### Determination of lesion volume

Mouse brains were cut into serial 20 µm cryostat sections. Every 10th section was stained with cresyl violet and the lesion area was determined using the Image J analysis software. Sections were digitized and converted to gray scale, and the border between TBI and non-TBI tissue was outlined. The lesion volume was calculated by subtracting the volume of the non-lesioned area in the ipsilateral hemisphere from the volume of the whole area in the contralateral hemisphere and expressed in mm^3^ as previously described [32].

### Assessment of edema

Brain water content is a sensitive measure of cerebral edema which was determined using the wet/dry method as previously described [32]. Mice were decapitated and brains were rapidly removed from the skull. The fresh brain was weighed in 3 mm coronal sections of the ipsilateral cortex, centered upon the impact site, then dehydrated for 24 h at 110°C and reweighed. The percentage of brain water content was calculated using the following formula: (wet weight - dry weight) / (wet weight) × 100.

### Tracer injection to detect BBB leakage

Mice were injected via the tail vein with Alexa Fluor 555-cadaverine (6 μg/g; Invitrogen, A30677) [33] dissolved in saline 1, 3 and 8 days after mTBI. After 2 h mice were anesthetized and perfused with phosphate-buffered saline (PBS) at pH 7.4, and the brains were collected.

### Evaluation of BBB permeability

BBB permeability was assessed by measuring the extravasation of Evans Blue dye as described previously [34]. 24 h after TBI injury, the mice were anesthetized, and Evans Blue dye (2% in saline) was injected slowly through the jugular vein (4 ml/kg) and allowed to circulate for 1.5 h before sacrifice.

### FAM-Aβ40 injection and tracing *in vivo*

In brief, 1 ng of human FAM-Aβ40 (Anaspec, # AS-23514-01) [35] in a total volume of 100 nl in saline was stereotaxically injected into ipsilateral hemisphere within the mTBI affected cortical region and unaffected contralateral hemisphere using the Neurostar motorized ultra-precise small animal stereotaxic instrument (Model 963SD). Aβ fluorescent signal distribution in the cortical regions 30 min after different injection was determined using high-resolution confocal microscopy.

### *in vivo* BBB efflux of Aβ40

CNS clearance of synthetic human FAM-Aβ40 peptides was determined in mice using a procedure we described previously [35]. Briefly, brain and blood were sampled 30 min after Aβ injection and prepared for Aβ40 ELISA. Simultaneously injection of ^14^C-inulin (reference marker) was also performed. The percentage of Aβ40 or inulin remaining in the brain after microinjection was determined as percentage recovery in brain = 100 × (Nb/Ni), where Nb is the amount of Aβ40 or ^14^C-inulin remaining in the brain at the end of the experiment and Ni is the amount of Aβ40 or ^14^C-inulin simultaneously injected into the brain ISF. The percentage of Aβ cleared through the BBB was calculated as [(1 - Nb(Aβ)/Ni(Aβ)) - (1 - Nb(inulin)/Ni(inulin))] × 100, using a standard time of 30 min [35].

### Laser Speckle Contrast Imaging (LSCI)

LSCI is based on the blurring of interference patterns of scattered laser light by the flow of blood cells to visualize blood perfusion in the microcirculation instantaneously [36]. Briefly, mice were anesthetized with gas anesthesia (isoflurane 2% in oxygen), with the head fixed in a stereotaxic frame (Kopf Instruments) and placed under an RFLSI Pro (RWD life sciences) and maintained at 1.2% isoflurane throughout the experiment. The surface of the skull is illuminated with a 784-nm 32-mW laser (RWD life science) at a 30-deg angle with a beam expander and light intensity controlled by a polarizer. Blood flow is detected by a CCD camera and the image acquisition is performed using custom software (RWD life science). Three hundred frames are acquired at 10 Hz with 10-ms exposure time. For the assessment of speckle contrast over time, regions of interest (ROIs) were selected and centered at the site where the laser is targeted on the cortex.

### Immunohistochemistry and confocal microscopy analysis

At endpoint, mice were anesthetized and transcardially perfused with PBS, and mouse brains were extracted and fixed for 12 h in 4% paraformaldehyde (PFA) in PBS at 4 °C. 20 µm-thick coronal cryosections were used for immunohistochemistry in this study. After washing with PBS, brain sections were permeabilized and incubated in 5% Donkey Serum for 1 h for blocking. Then the brain tissues were incubated in primary antibody overnight at 4 °C. The primary antibody information is as following: anti-Fibrinogen antibody (1:200, Dako, A0080), anti-Iba1 antibody (1:500, Wako, 019-19741), anti-NeuN antibody (1:200, Abcam, ab177487), anti-Glial fibrillary acidic protein (GFAP) antibody (1:200, Invitrogen, 13-0300), anti-SMI-32 antibody (1:1000, Biolegend, 801709), anti-Cluster of differentiation (CD13) antibody (1:200, R&D Systems, AF2335), anti-β amyloid antibody (1:200, #D54D2, Cell Signaling). After washing off the primary antibody with PBS, the brain sections were then incubated with secondary antibodies (1:500; Jackson Immunoresearch Laboratories) for 1 h at room temperature. Dylight 488 conjugated-lectin (1:200, Vector, DL1174) was used to label brain vasculature, and Alexa Fluor 647 Donkey anti-Mouse IgG (1:500; Invitrogen, A-31571) was used to detect extravasation of IgG. After that, the sections were rinsed with PBS and covered with fluorescence mounting medium with DAPI (4’,6-diamidino-2-phenylindole; Vector Laboratories, #H-1200).

### Detection of apoptosis

TUNEL assay was performed according to the manufacturer’s protocol (CAS7791-13-1, Roche). Tissues were incubated with the TUNEL reaction mixture in a humidified chamber for 30 min at 37 °C in the dark.

### Detection of Aβ plaques

After PFA fixation, the sections were incubated in 1% aqueous Thioflavin S (T1892; Sigma) for 5 min and rinsed in 80% ethanol, 95% ethanol, and distilled water. All images were taken with the Nikon A1R confocal microscopy and analyzed using NIH ImageJ software. In each animal, 4 randomly selected fields in the cortex and hippocampus in TBI-affected hemisphere were analyzed in 4 non-adjacent sections (∼100 µm apart) and averaged per mouse.

### Hematoxylin staining

The brain tissue sections were completely covered with adequate Hematoxylin (Vector, H-3502) and incubated for 5 min; next, the slides were rinsed in 2 changes of distilled water (15 seconds each) to remove the excess stain; then the slides were rinsed using 100% ethanol (10 seconds). At last, the slides were coverslipped with mounting medium.

### Extravascular IgG, fibrinogen and fibrin deposits

We used antibodies that detected IgG, fibrinogen and fibrinogen-derived fibrin polymers (see Immunohistochemistry). Ten-micron maximum projection z-stacks were reconstructed, and the IgG, fibrinogen and fibrin-positive perivascular signals on the abluminal side of lectin-positive endothelial profiles on microvessels ≤ 6 µm in diameter were analyzed using ImageJ. In each animal, 4-5 randomly selected fields in the cortex in both mTBI-affected hemisphere and unaffected contralateral hemisphere were analyzed in 4 non-adjacent sections (∼100 µm apart) and averaged per mouse.

### Analysis for SMI-32

Ten-micron maximum projection z-stacks were reconstructed, and the SMI-32 signals were analyzed using ImageJ. In each animal, 4 randomly selected fields in the cortex and hippocampus in TBI-affected hemisphere were analyzed in 4 non-adjacent sections (∼100 µm apart) and averaged per mouse.

### GFAP-positive astrocytes and Iba1-positive microglia counting

For quantification, GFAP-positive astrocytes or Iba1-positive microglia that also co-localized with DAPI-positive nuclei were quantified by using the Image J Cell Counter analysis tool. In each animal 5 randomly selected fields from the cortex were analyzed in 4 nonadjacent sections (∼100 µm apart). The number of GFAP-positive astrocytes was expressed per mm^2^ of brain tissue. The activated microglia undergo morphological changes including retraction of the processes and acquisition of a phagocytotic shape, but the quiescent microglia exhibit a ramified morphology [37].

### NeuN-positive neuronal nuclei counting

For quantification, NeuN-positive neurons were quantified by using the Image J Cell Counter analysis tool. In each animal, 5 randomly selected fields from the cortex were analyzed in 4 nonadjacent sections (∼100 µm apart). The number of NeuN-positive neurons colocalized with TUNEL was counted followed by the same procedure.

### Quantification of pericyte numbers and coverage

The quantification analysis of pericyte numbers and coverage was restricted to CD13-positive perivascular mural cells that were associated with brain capillaries defined as vessels with ≤ 6 µm in diameter. For pericyte numbers, ten-micron maximum projection z-stacks were reconstructed, and the number of CD13-positive perivascular cell bodies that co-localized with DAPI-positive nuclei on the abluminal side of lectin-positive endothelium on vessels ≤ 6 µm counted using ImageJ Cell Counter plug-in. In each animal, 4-5 randomly selected fields (640 × 480 µm) in the cortex regions were analyzed in 4 non-adjacent sections (∼100 µm apart) and averaged per mouse. The number of pericytes was expressed per mm^2^ of tissue. For pericyte coverage, ten-micron maximum projection z-stacks (area 640 × 480 µm) were reconstructed, and the areas occupied by CD13-positive (pericyte) and lectin-positive (endothelium) fluorescent signals on vessels ≤ 6 µm were subjected separately to threshold processing and analyzed using ImageJ as previously described [38]. In each animal, 4-5 randomly selected fields in the cortex regions were analyzed in 4 nonadjacent sections (∼ 100 µm apart) and averaged per mouse.

### Microvascular capillary length

Ten-micron maximum projection z-stacks were reconstructed, and the length of lectin-positive capillary profiles (≤ 6 µm in diameter) was measured using the ImageJ plugin “Neuro J” length analysis tool. In each animal, 4-5 randomly selected fields (640 × 480 µm) in the cortex were analyzed from 4 nonadjacent sections (∼100 µm apart) and averaged per mouse. The length was expressed in mm of lectin-positive vascular profiles per mm^3^ of brain tissue.

### Aβ deposits calculation

Aβ-positive areas were determined using ImageJ software. Briefly, the images were taken on a BZ9000 fluorescent microscope in single plain at 20× (640 × 480 µm image size) and subjected to threshold processing (Otsu) using ImageJ, and the percent area occupied by the signal in the image area was measured as described previously [31]. In each animal, four randomly selected fields from the cortex and hippocampus were imaged and analyzed in four nonadjacent sections (∼ 100 µm apart).

### Western blot

The total proteins from cortical tissue of TBI-affected hemisphere in sham-operated, sTBI and mTBI mice were extracted, and Synapsin I was analyzed by Western blot. Briefly, protein samples were separated by electrophoresis on a 10% polyacrylamide gel and electrotransferred to a nitrocellulose membrane. Nonspecific binding sites were blocked in TBS, overnight at 4°C, with 2% BSA and 0.1% Tween-20. Membranes were rinsed for 10 min in a buffer (0.1% Tween-20 in TBS) and then incubated with anti-synapsin I (1:1500, Sigma) followed by anti-rabbit IgG horseradish peroxidase conjugate (Santa Cruz Biotechnology). After rinsing with buffer, the immunocomplexes were developed with G:BOX Chemi XX6 gel doc system (Syngene) and analyzed in Image Lab software. β-Actin was used as an internal control for Western blot.

### Statistical analysis

Sample sizes were calculated using nQUERY assuming a two-sided alpha-level of 0.05, 80% power, and homogeneous variances for the 2 samples to be compared, with the means and common standard deviation for different parameters predicted from published data and pilot studies. For comparison between two groups, the F test was conducted to determine the similarity in the variances between the groups that are statistically compared, and statistical significance was analyzed by Student’s t-test. For multiple comparisons, Bartlett’s test for equal variances was used to determine the variances between the multiple groups and one-way analysis of variance (ANOVA) followed by Bonferroni’s post hoc test was used to test statistical significance. All analyses were performed using GraphPad Prism 8 software by an investigator blinded to the experimental conditions. A *p*-value < 0.05 was considered as statistically significant.

## Results

### Microvascular injury in a mouse model of mTBI

To determine the vascular impairment in mTBI, we first compared an open-head sTBI model with a close-head mTBI model based on controlled cortical impact (**Sup Fig. 1**). Consistent with previous studies [7], the sTBI model induced strong deficits in motor function, as seen on both rotarod test and foot-fault test (**Fig. 1A-B**); whereas the mTBI model only induced transient motor impairments, as the mice nearly recovered in 3 days in the same behavioral tests (**Fig. 1A-B**). In line with the behavioral outcomes, mTBI model didn’t cause visible cortical lesion at 24 h post-injury (**Fig. 1C-D**), compared to the sTBI model with an average lesion volume of over 2 mm^3^ (**Fig. 1C-D**). Brain water content, a sensitive measure of cerebral edema, was not significantly changed 24 h after mTBI either (**Sup Fig. 2A**). In addition, we observed very few TUNEL- and NeuN-double-positive cells in the mTBI impacted ipsilateral cortex at 3 days after mTBI (**Sup Fig. 2B-C**); while ∼20% neuronal death was found near the cortical lesion in sTBI model (**Sup Fig. 2B-C**). Therefore, the data suggests that our murine mTBI model is analogous to concussion [39].

**Fig. 1.**
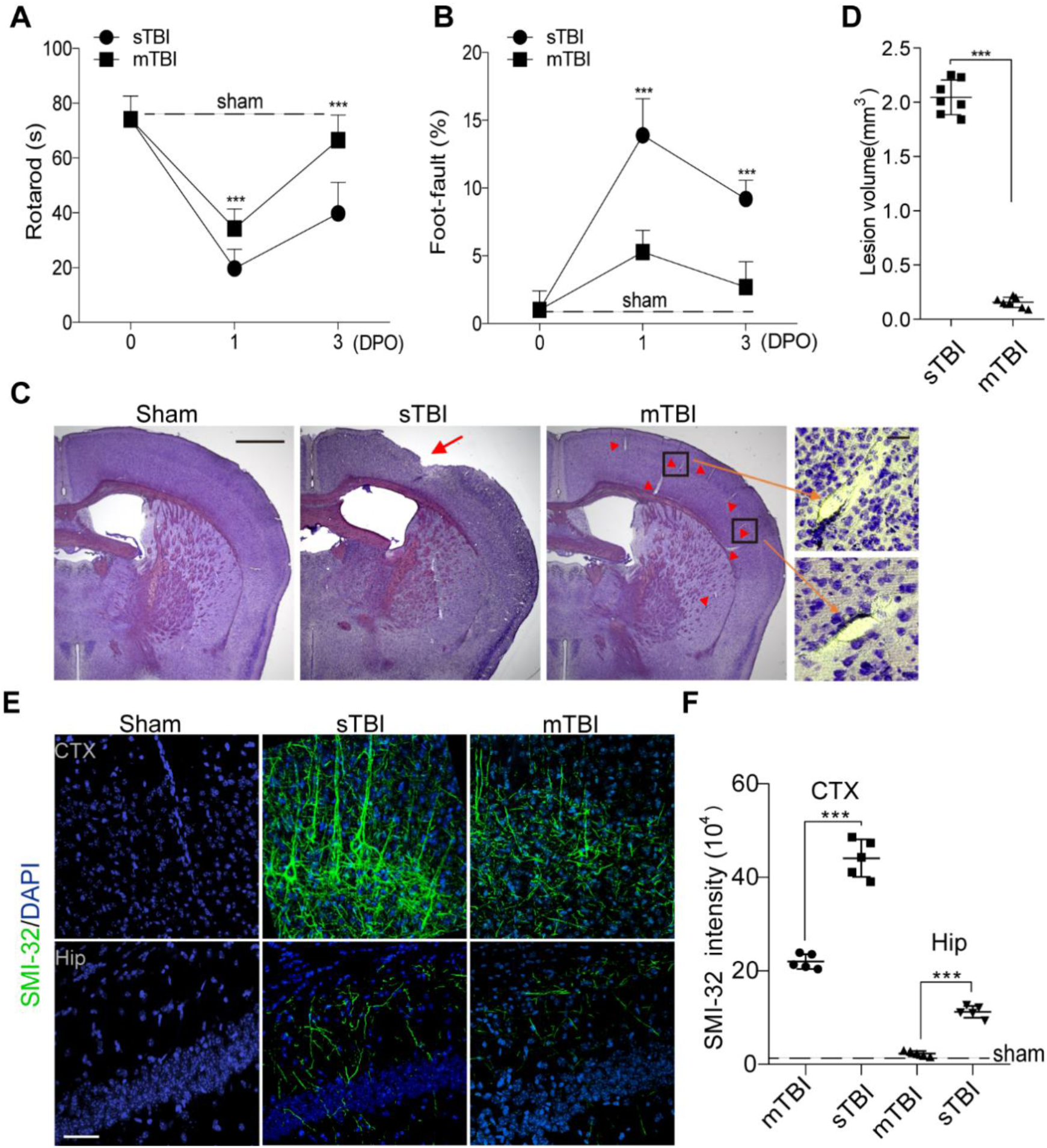
Traumatic brain injury models with controlled cortical impact in mice. (**A-B**) Rotarod test (**A**) and foot-fault test (**B**) were performed on 0, 1 day and 3 days post operation (DPO) in sTBI mice (*n = 8*) or mTBI mice (*n = 7*). Dash lines indicate average value from sham-operated group (*n = 5*). (**C**) Representative images of hematoxylin staining of brain sections from sham-operated, sTBI and mTBI mice. Scale bar = 1 mm (left), and 50 μm (right) in high magnification. Arrow indicates cortical lesion; arrowhead indicates enlarged perivascular space. (**D**) Quantification of TBI-induced lesion volume at 3 days post-injury in sTBI mice and mTBI mice. The injury resulted in a significantly larger lesion volume in sTBI mice when compared with mTBI mice. *n = 7* mice per group. (**E-F**) Representative images (**E**) of fluorescence immunostaining for SMI-32 (green) and DAPI (blue), quantification of SIM-32 fluorescence intensity (**F**) in cortex (CTX) and hippocampus (Hip) of sham-operated, sTBI and mTBI mice 3 days after injury. Scale bar = 50 μm, *n = 5* mice in sham-operated, sTBI and mTBI group respectively. In (**A-B**), data are presented as Mean ± SD; ***, *P* < 0.001; NS, non-significant (*P* > 0.05), one-way ANOVA followed by Bonferroni’s post-hoc tests. In **D, F**, Mean ± SD; ***, *P < 0.001* by Student’s t-test.

Interestingly, axonal injuries were detectable 3 days after mTBI in both mTBI-affected ipsilateral cortex and hippocampus (**Fig. 1E**), based on a monoclonal SMI-32 antibody against non-phosphorylated neurofilament [40], albeit to a lesser extent than that in sTBI mice (**Fig. 1E-F**). Western blotting of the synaptic marker, synapsin I, also indicated a ∼32% reduction in synaptic proteins in mTBI-affected ipsilateral cortex, compared to a 75% reduction after sTBI (**Sup Fig. 2D-E**). Microglial activation and astrogliosis were also substantially reduced in both mTBI-affected ipsilateral cortical areas and contralateral area when compared with sTBI-affected ones, as indicated by Iba1 and GFAP immunostainings (**Sup Fig. 3**). For example, nearly 90% Iba1-positive microglia morphologically resemble the activated phagocytic form in sTBI-affected ipsilateral area, while ∼51% of them remained in ramified resting morphology [41] after mTBI (**Sup Fig. 3B**). On the other hand, reactive GFAP-positive astrocyte numbers were also much lower in mTBI, when compared with the same hemispheric cortical regions in sTBI (**Sup Fig. 3C-D**). Unlike sTBI, the microglial activation and astrogliosis did not spread much to the contralateral hemisphere at 3 days after mTBI (**Sup Fig. 3B, D**).

BBB breakdown is a hallmark of TBI, which occurs acutely within hours after tissue damage and lasts for days [42]. As hematoxylin staining showed vascular segments accompanied by enlarged perivascular space in brain sections from mTBI mice (**Fig. 1C**, arrowheads and inserts), indicating vascular impairment, we next performed histological analysis to examine the microvascular injury and BBB dysfunction occurred in the acute/subacute phase of mTBI. Immunostaining for plasma-derived IgG (**Fig. 2A**) demonstrated a ∼6-fold increase in perivascular IgG deposits in the mTBI impacted cortical area 3 days after mTBI (**Fig. 2B**). We also found similar results in extravascular fibrin deposits (**Sup Fig. 4A, B**), or extravasation of exogenous tracer cadaverine [33] (**Sup Fig. 4C, D**), suggesting BBB is compromised after mTBI, which is consistent with findings in human patients and different animal models [43].

**Fig. 2.**
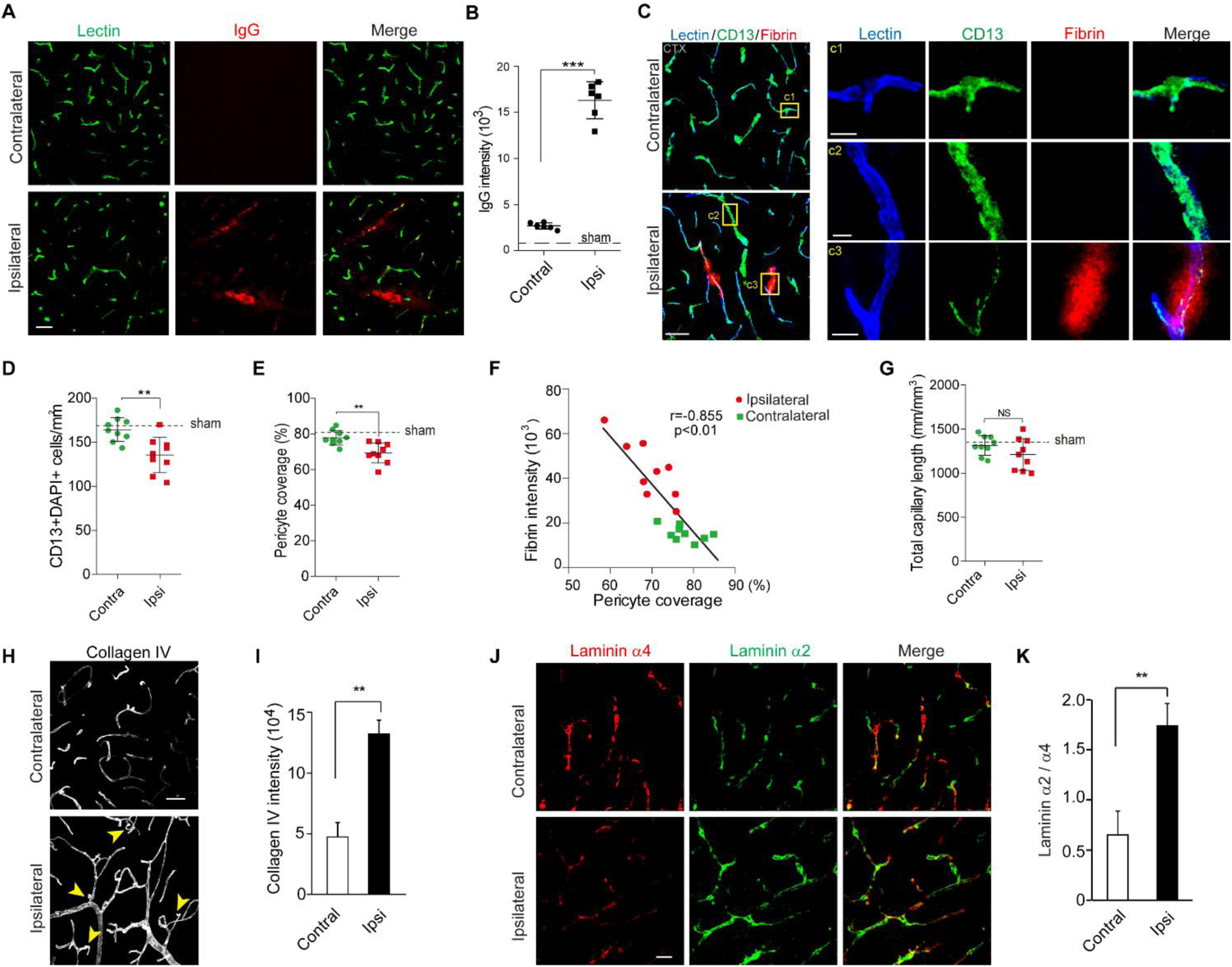
Altered pericyte coverage and basement membrane structure in mTBI mice. (**A-B**) Representative images (**A**) and quantification (**B**) showing extravasation of plasma IgG (red) in brain parenchyma 3 days after mTBI. n = 6 mice. Dash line indicates an average value from sham-operated group (*n = 5* mice). Scale bar = 50 µm. (**C**) Representative images showing CD13-positive pericyte coverage (green), lectin-positive endothelial vascular profiles (blue) and extravasation of fibrin deposits (red) in brain parenchyma of mTBI-affected ipsilateral side or contralateral side 3 days after mTBI. Scale bar = 50 µm. Insert on right: high magnification images of boxed regions in (c1-c3), Scale bar = 10 µm. (**D-E**) Quantification of CD13-positive pericyte cell bodies per mm^2^ (**D**) and pericyte coverage on lectin microvessel profiles (**E**) in brain parenchyma of mTBI-affected ipsilateral side and contralateral side 3 days after mTBI. *n = 9* mice per group. (**F**) Pearson’s correlation plot between BBB impairment based on fibrin deposits and loss of pericyte coverage in brain parenchyma 3 days after mTBI. *n* = *total 18* hemispheres. r = Pearson’s coefficient. *P*, significance. (**G**) quantification of microvascular density in the cortexes of mTBI-affected ipsilateral side and contralateral side 3 days after mTBI. *n = 9* mice per group. (**H-I**) Representative confocal microscope images (**H**) and quantification (**I**) showing upregulation of collagen IV in mTBI-impacted ipsilateral cortical area 3 days after mTBI. Scale bar = 50 µm. (**J-K**) Representative confocal microscope images (**J**) and quantification (**K**) showing laminin α2 and α4 changes in mTBI-impacted ipsilateral cortical area 3 days after mTBI. Scale bar = 30 µm. *n = 6* mice per group. Mean ± SD in **B, D-E, G, I, K**; **, *P* < 0.01; ***, *P* < 0.001; NS, non-significant (*P* > 0.05) by Student’s t-test.

### mTBI resulted in pericytes loss and basement membrane alteration

Pericytes play a key role in regulating neurovascular functions of the brain including formation and maintenance of the BBB [33]. To determine whether mTBI influences pericytes, we next performed confocal imaging analysis for CD13-positive pericytes [33] (**Fig. 2C**). We found a substantial loss of brain pericytes based on the number of CD13-positive cells (**Fig. 2D**) in the impacted ipsilateral side 3 days after mTBI when compared with unaffected contralateral hemisphere, as well as a reduction of pericyte coverage based on CD13 profile on Lectin-positive endothelial profiles [38] (**Fig. 2E**), which is even more evident in microvascular segment with strong fibrin deposits (**Fig. 2C** inserts). More importantly, accumulation of perivascular fibrin deposits in mTBI mice correlated tightly with the reductions in pericyte coverage (**Fig. 2F**). However, there were no significant changes in total capillary length between impacted ipsilateral hemisphere and unaffected contralateral hemisphere (**Fig. 2G**), suggesting acute vascular rarefication perhaps is more specific to moderate and severe TBI [44].

Recent genetic profiling studies in rodents using microarray [45] and RNA sequencing [46] indicated a TBI-induced genomic response in genes associated with the extracellular matrix proteins and basement membrane structures, which are in line with our observation of enlarged perivascular space and pericyte loss. We then performed histological analysis for different markers of perivascular compartments and extracellular basement membrane. Collagen IV, secreted by endothelial cells, mural cells and astrocytes, is a marker for both inner and outer basement membranes in the perivascular space [47]. It was significantly increased by 2.9-fold in mTBI-affected hemisphere compared to the unaffected contralateral side of the brain (**Fig. 2H-I**). More interestingly, we found immunostaining for vascular laminin α4 [48] was dramatically reduced in the mTBI-affected hemisphere when compared with the unaffected contralateral hemisphere (**Fig. 2J**, left), which is consistent with vascular injury and suggests vascular compartment within the perivascular space decreased after mTBI. In addition, laminin α2 immunoreactivity around the astrocytic endfeet of microvessels [49] was dramatically increased in mTBI-affected hemisphere compared to the unaffected contralateral hemisphere (**Fig. 2J**, middle), indicating that the parenchymal compartment of the perivascular space increased after mTBI. Therefore, the ratio of laminins α2/α4 may potentially reflect the structural changes of basement membrane and perivascular space after mTBI (**Fig. 2K**). Furthermore, the pathological changes of BBB in the acute/subacute phase of mTBI was associated with microcirculation insufficiency, as LSCI showed a ∼50% reduction in CBF in the impacted cortex 3 days after mTBI (**Sup Fig. 4E, F**).

### mTBI induced BBB dysfunction precedes gliosis

To map the BBB dysfunctions in the acute/subacute phase of mTBI, we performed tracing of Evans blue at 1 day after impact (**Fig. 3A**). The breakdown of the BBB mainly occurred right underneath the impact area, consisting primarily the somatosensory and motor cortexes (**Fig. 3A**), with the upper layers of cortex subjected to accumulation of Evans blue after mTBI (**Fig. 3B-C**). The penetration of vascular damage correlated with the physical impact; as BBB leakage peaked around the impact center (**Fig. 3D-E**). On the other hand, sTBI produced not only substantial tissue lesion in the cortex (**Sup Fig. 5A-C**), but also resulted in a much larger area with a leaky BBB to Evans blue (**Sup Fig. 5A-C**), which often penetrated to hippocampal regions (**Sup Fig. 5C**), as indicated by quantification of BBB leakage and tissue damage (**Sup Fig. 5D-E**).

**Fig. 3.**
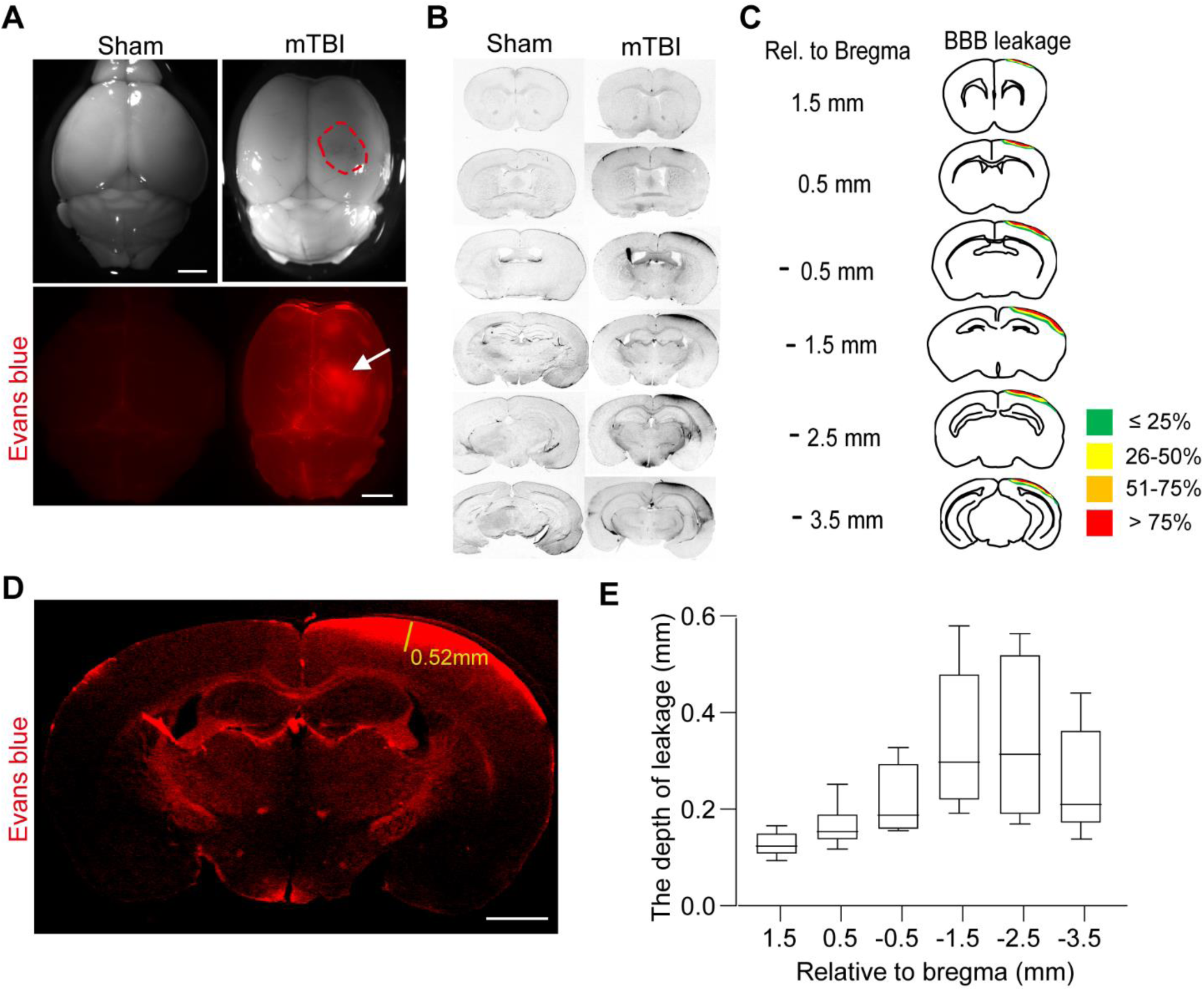
Mapping BBB dysfunction in mTBI mice. (**A**) Brain accumulation of Evans blue was detected at 1 day post injury under both brightfield microscope and fluorescent microscope, indicating the disruption of BBB in mTBI mice. Red dashed circle and white arrow indicate the area of BBB leakage. Scale bar = 1 mm. (**B**) Representative grey scale images showing Evans blue in coronal brain sections at every 1 mm relative to bregma from mice in (A). (**C**) Schematic accumulative map of the BBB disruption based on Evans blue leakage at 1 day after mTBI. Each slide depicts the accumulated information of BBB distribution (color coded as indicated) at the given section in relation to the bregma. *n = 10* mice. (**D-E**) Representative coronal brain section from an mTBI mice showing BBB leakage in brain parenchyma 1 day after mTBI (**D**) and the whisker plot quantification (**E**) for the depth of BBB leakage at each section position. Scale bar = 1 mm, *n = 10* mice. Data are presented in box plots showing the upper (75%) and lower (25%) quartiles, with the horizontal line as the median and the whiskers as the maximum and minimum values observed.

To examine the time course of cerebrovascular impairment and BBB dysfunction in mTBI mice, we also performed *in vivo* tracing of systemically administrated Alexa Fluor 555-cadaverine [33], on 1 day, 3-and 8-days post operation (DPO). Using the confocal microscopy imaging (**Fig. 4A**), we found that a severe BBB impairment in the impacted cortex area at 1 DPO as indicated by a ∼30.1-fold increase of parenchymal cadaverine uptake, which recovered gradually around 3 and 8 DPO (**Fig. 4C**). On the other hand, the BBB impairment in the hippocampal area appeared at 1 DPO, but peaked around 3 DPO as indicated by ∼24.1-fold and ∼20.7-fold increase of uptake in CA3 and dentate gyrus (DG) areas, respectively (**Fig. 4D-E**).

**Fig. 4.**
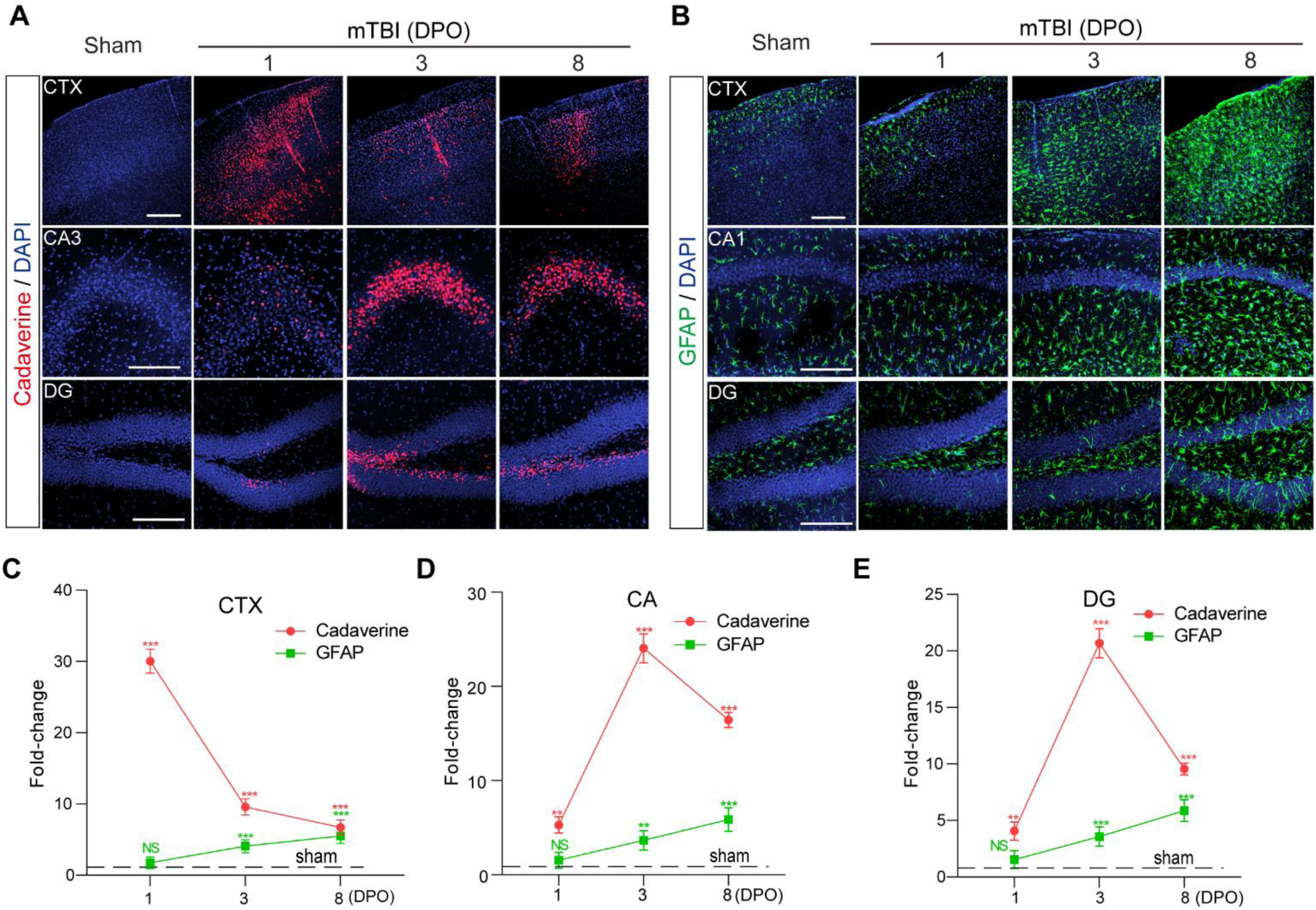
The time courses of BBB dysfunction and astrogliosis after mTBI in mice. (**A**) Representative confocal microscope images showing the extravasation of intravenously administrated Alexa-555 cadaverine (red) in cortex (CTX), Cornu Ammonis 3 (CA3) and dentate gyrus (DG) area 1-day, 3- and 8-days post operation (DPO). Scale bar = 100 µm. (**B**) Representative confocal microscope images showing GFAP-positive astrocytes (green) in cortex (CTX), CA1 and dentate gyrus (DG) area 1-day, 3- and 8-days post operation (DPO). Scale bar = 100 µm. (**C-E**) Quantification for the fold changes for cadaverine intensity (*n = 5* mice per time point) and GFAP positive cells (*n = 5* mice per time point) in CTX area, CA area and DG area 1-day, 3- and 8-days post operation (DPO). In **C-E**, data are presented as mean ± SD; ***, *P* < 0.001; **, *P* < 0.01 NS, non-significant (*P* > 0.05), one-way ANOVA followed by Bonferroni’s post-hoc tests. Dash lines indicate the sham-operated group.

Next, we examined the time course of astrogliosis and microglial activation after mTBI. Immunostaining for GFAP-positive astrocytes (**Fig. 4B**) and Iba1-positive microglial cells (**Sup Fig. 6A**) indicated that astrocytes and microglial cells were substantially activated on 3 DPO and keep raising gradually to 8 DPO after mTBI in both cortex and hippocampus (CA1 and DG) areas when compared to the number of astrocytes and microglial cells in sham-operated mice (**Fig. 4C-E** and **Sup Fig. 6B-D**). Therefore, our data indicate that microvascular injury and BBB dysfunction preceded astrogliosis and microglial activation in the acute/subacute phase of mTBI.

### mTBI exacerbated amyloid pathologies and cognitive impairment in 5xFAD mice

Prominent amyloid pathologies are found in at least 30% of TBI patients [47]. This direct link between TBI and amyloid pathologies has been partially attributed to increased Aβ production [5]. However, it is currently unknown whether TBI impairs Aβ clearance via the vascular system, and if so, to what extent TBI-induced vascular impairment exacerbated amyloid pathologies and cognitive impairment *in vivo*. To address these questions, we conducted mTBI in the 5xFAD mouse model of AD at 3 months of age, where amyloid pathology is evident but cognitive function remains relatively normal [26].

Longitudinal behavioral tests in the acute/subacute phase of mTBI indicated that the 5xFAD mice had a transient impairment in motor coordination at 1 DPO but soon recovered nearly back to normal in 8 days, as shown by rotarod and foot fault tests (**Fig. 5A-B**). However, these 5xFAD mice suffered substantially from cognitive impairment at 8 DPO, as shown by >60% reduction in freezing time based on contextual fear conditioning test when compared with sham-operated 5xFAD mice or wild type littermates (WT) (**Fig. 5C**). mTBI also exacerbated the BBB dysfunction in 5xFAD mice [31], as shown by >53.3% extravascular fibrin deposits in both mTBI affected cortex and hippocampus when compared with sham-operated mice at 8 DPO (**Fig. 5D, E**). Additionally, mTBI resulted in gliosis changes in cortex and hippocampus, as indicated by significant increases in both astrocytes (**Fig. 5F, G**) and activated microglia cells at 8 DPO (**Fig. 5H, I**).

**Fig. 5.**
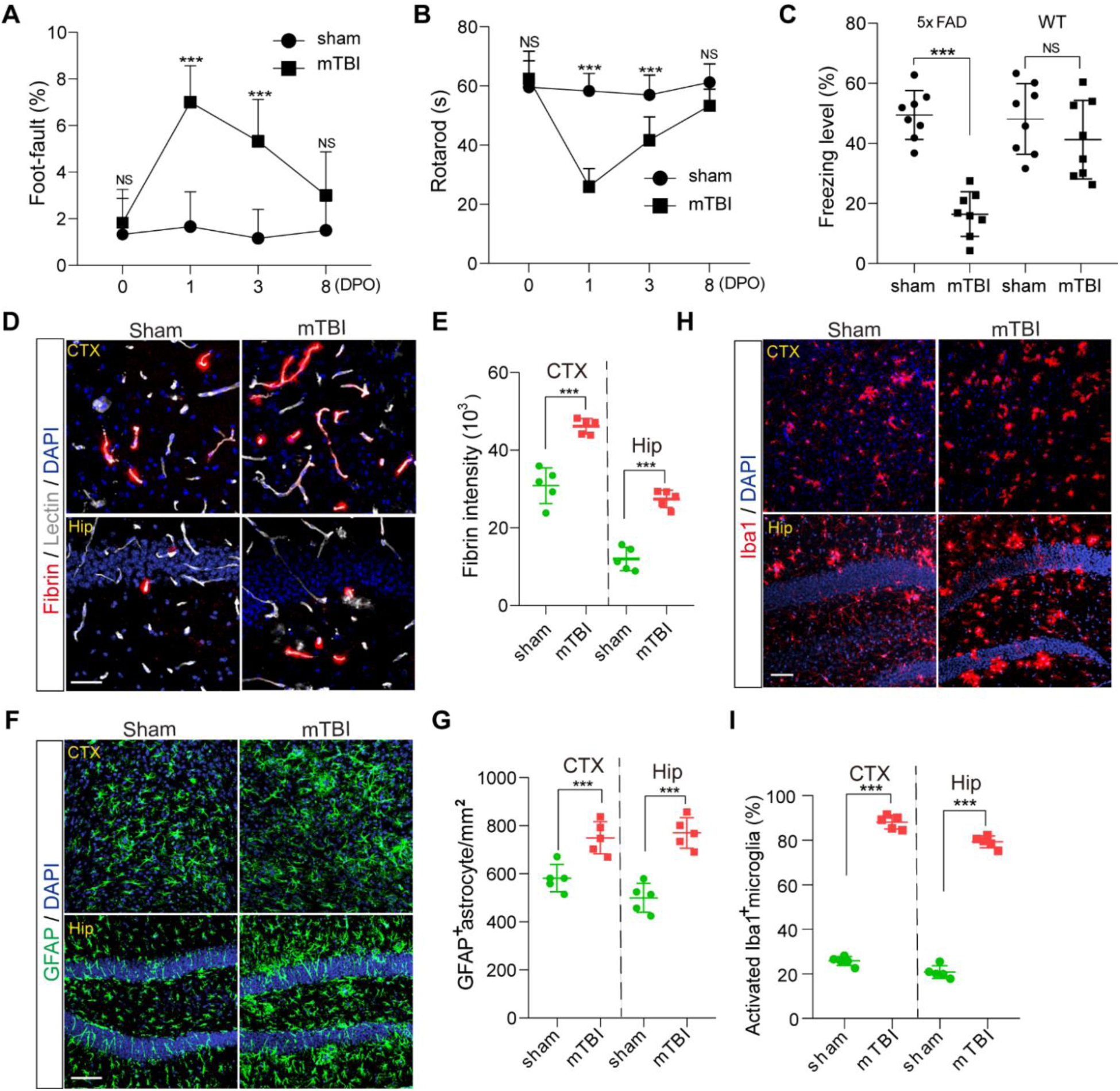
Accelerated cognitive impairment and pathologies in 5xFAD mice after mTBI. (**A-B**) Foot-fault (**A**) and Rotarod (**B**) tests were performed in 3 months old 5xFAD mice on 0, 1, 3 and 8 days after mTBI or sham-operation. *n = 8* mice per time point per group. (**C**) Contextual fear conditioning test was performed in 3 months old 5xFAD mice 8 days after mTBI or sham-operation as in (**A-B**), or in non-transgenic littermates (WT) 8 days after mTBI or sham-operation. *n = 8* mice per group. (**D-E**) Representative images (**D**) and quantification (**E**) showing extravascular fibrin deposits in brain parenchyma 8 days after mTBI, in both cortex (CTX) and hippocampus (Hip). Lectin (gray), fibrin (red). *n = 5* mice; Scale bar = 50 µm. (**F-G**) Representative images (**F**) and quantification (**G**) showing GFAP-positive astrocytes in brain parenchyma 8 days after mTBI. GFAP (green). *n = 5* mice per group; Scale bar = 50 µm. (**H**) Representative images of fluorescence immunostaining for microglial marker Iba1 (red) and DAPI (blue) in brain parenchyma 8 days after mTBI. Scale bar = 50 µm. (**I**) Quantification for the percentage of activated Iba1 positive cells in both cortex and hippocampus (*n = 5* mice per group). In **A-B**, Mean ± SD; ***, *P* < 0.001, NS, non-significant (P > 0.05), one-way ANOVA followed by Bonferroni’s post-hoc tests. In **C, E, G, I**, Mean ± SD; ***, *P* < 0.001, NS, non-significant (*P* > 0.05) by Student’s t-test.

More importantly, using histological analysis with rabbit monoclonal antibody for a broad range of Aβ species (**Fig. 6A**), we found that the amyloid burden in mTBI mice was nearly doubled in both cortex and hippocampus 8 days after mTBI when compared with sham-operated 5xFAD mice (**Fig. 6B**). This is consistent with increased Thioflavin-S positive amyloid plaques (**Fig. 6C-D**), and demonstrates that mTBI exacerbated the amyloid pathologies in 5xFAD mice.

**Fig. 6.**
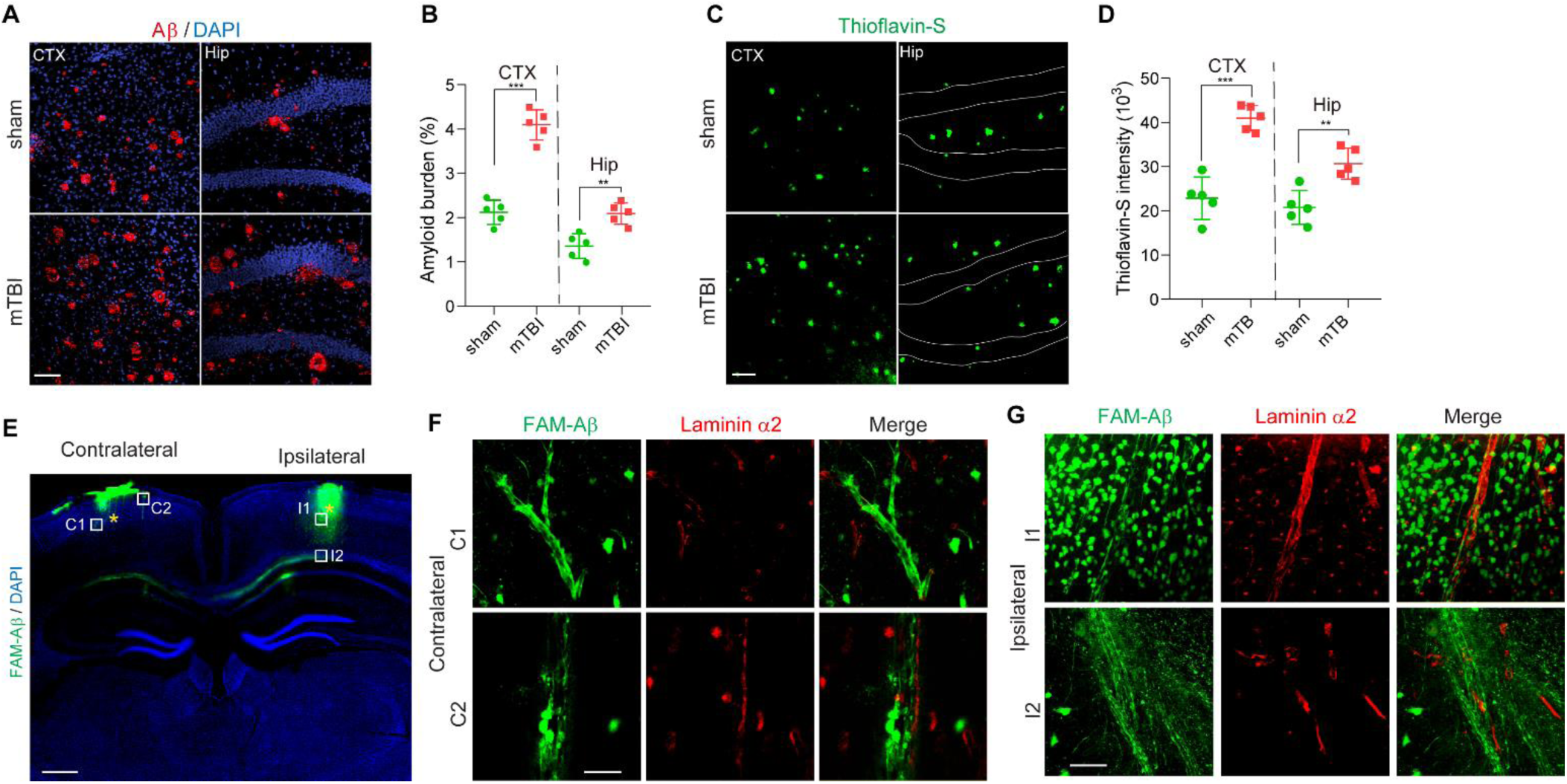
Accelerated amyloid pathology in 5xFAD mice after mTBI. (**A-B**) Representative images (**A**) showing immunostaining of brain amyloid deposits (red), DAPI (blue) and quantification (**B**) for the amyloid burden (based on the percentage of cortical area occupied with Aβ immunostaining) in cortex (CTX) and Hippocampus (Hip) area 8 days post operation. *n = 5* mice per group, scale bar = 100 µm. (**C-D**) Representative images (**C**) showing Thioflavin-S labelled amyloid plaques (green) and quantification for the intensity of Thioflavin-S (**D**) in cortex (CTX) and Hippocampus (Hip) area 8 days post operation. *n = 5* mice per group, scale bar = 100 µm. In **B** and **D**, data are presented as Mean ± SD; ***, *P* < 0.001; **, *P* < 0.01; by Student’s t-test. (**E**) Representative tiled confocal images showing distribution of FAM-Aβ (green) in a coronal section, 30 min after bilateral injection in mTBI mice. Bar = 0.5 mm. Asterisks indicate the injection sites. DAPI: nuclei staining. (**F-G**) High magnification images of boxed regions in contralateral side (**F**, C1-C2) and mTBI-impacted ipsilateral side (**G**, I1-I2), showing differential distribution of FAM-Aβ40. Laminin α2: basement membrane marker. Scale bar = 20 µm.

### mTBI impaired brain Aβ clearance through the vascular pathways

More recent studies in mice using fluorescent solutes demonstrated a brain clearance system drains interstitial fluid (ISF) and macromolecules along the vasculature and eventually connects to the lymphatic system in the dura mater, for waste disposal to the peripheral immune system [50]. More specifically, when fluorescently labeled Aβ was injected in the brain parenchyma, it followed the ISF flow, reaches the perivascular space and is passively eliminated through drainage [51]. Therefore, we next performed *in vivo* tracing of FAM (fluorescein) labeled human Aβ peptides, where 1 ng FAM-Aβ40 (Anaspec, # AS-23514-01) [35] was simultaneously injected into both ipsilateral and contralateral cortexes (**Sup Fig. 7A**). We determined Aβ fluorescent signal distribution in the cortical regions 30 min after different injections, and found that FAM-Aβ40 in the unaffected contralateral hemisphere was mainly distributed along penetrating vessels and transported back to the brain surface (**Fig. 6E**, contralateral). However, FAM-Aβ40 was retained in the parenchyma in mTBI affected hemisphere, with visible transport along axonal tracks that even reached cortico-callosal projections in the corpus callosum (**Fig. 6E**, ipsilateral). High-resolution confocal imaging and 3D reconstruction further demonstrated that FAM-Aβ40 distributed along with the perivascular space in the unaffected contralateral hemisphere (**Fig. 6F**), while it was taken up by neuron-like cells in the cortex and diffused along the axons (**Fig. 6G**). The appearance of axonal trafficking of injected FAM-Aβ40 is consistent with intra-axonal Aβ pathology seen in patients, indicating that the shift from vascular clearance of Aβ to neuronal uptake may contribute to axonal damage and neuronal dysfunction after mTBI [7].

In addition, measurement of BBB efflux of injected synthetic Aβ (**Sup Fig. 7A**) using sensitive ELISA assays [35] also demonstrated that brain retention of Aβ in mice was significantly increased at 3 days after mTBI (**Sup Fig. 7B**), whereas the amount of synthetic Aβ being transported across the BBB to the circulation was reduced more than 50% (**Sup Fig. 7C**). On the other hand, brain retention of inulin, an inert extracellular space tracer for estimating brain interstitial fluid to cerebrospinal fluid bulk flow [35], was increased (**Sup Fig. 7D**), suggesting a global inhibition of the perivascular drainage occurred after mTBI. Taken together, our data demonstrated that mTBI can induce vascular impairment and block brain clearance of Aβ through both perivascular drainage pathway and transvascular pathway, which contribute to the loss of Aβ homeostasis and subsequent Aβ accumulation in the brain of 5xFAD mice.

## Discussion

Microvascular injury and BBB impairment are commonly found across neurodegenerative conditions, including but not limited to TBI, stroke, AD, amyotrophic lateral sclerosis (ALS), multiple sclerosis (MS) and even aging [11,25], although the etiology in each condition could be rather distinct [11]. Vascular impairments induced by mTBI have been well documented both clinically in patients and preclinically in animal models, but the underlying mechanism has not been clearly understood. In the present study, we first established a rodent mTBI model, which was analogous to concussion in humans, as indicated by temporal locomotive behavioral deficits, and the lack of significant neuronal death or cortical tissue damage when compared with the sTBI model. We noted enlarged perivascular spaces surrounding blood vessels ipsilateral to mTBI as well as a significant increase in perivascular extravasation of plasma proteins and reduced CBF. These data are indicative of a compromised cerebral microvasculature, as well as a leaky and malfunctioning BBB.

With widening of the perivascular space and shrinking of the vascular compartment, perivascular drainage pathway became a victim of mTBI. We saw that synthetic FAM-Aβ40 retained in the mTBI-affected ipsilateral cortical area much more than in the contralateral side, which was accompanied by increased neuronal uptake and subsequent axonal trafficking of injected FAM-Aβ40 after TBI. We then used the 5xFAD mouse model of AD to further clarify the link between mTBI and AD, and found mTBI exacerbated cognitive deficits based on fear conditioning test. In conclusion, our study demonstrated that mTBI induces BBB dysfunction, damages the perivascular basement membrane, impairs the clearance of Aβ through perivascular drainage pathway, and exacerbates pathologies and cognitive deficiencies associated with AD.

It is reported that mTBI induces relatively sustained shear stress located within a discrete region near the impact contact zone [43], which contributes to acute and/or subacute brain pathologies, including microvascular injury and BBB disruption [6]. Subsequently, mTBI can elicit a complex sequence of cellular cascades, and lead to genomic responses and transcriptional changes in vascular cells [52], including vascular endothelial growth factor A (VEGFA) [53], and compensatory changes in the extracellular matrix proteins and expansion of the basal lamina [24, 44], which not only alter the basement membrane structure and perivascular space but also affect brain infiltration of peripheral leukocytes. Recent evidence from RNA sequencing of brain endothelial cells in different animal models has demonstrated that similar vascular gene expression profiles exist among multiple neurodegenerative diseases including TBI [54], suggesting perhaps that the molecular signature of vascular impairment might be common. Nevertheless, future studies exploring vascular associated transcriptomic changes at single-cell level in both mouse models and human patients are necessary for defining the exact microvascular molecular signature in mTBI, and beyond.

There has been growing attention to mTBI and the development of dementia later in life [55]. Potential underlying mechanisms between TBI and later development of AD include diffused axonal damage, persistent inflammation and vascular dysfunctions [7,56]. While aberrant Tau pathology induced by TBI is considered a high-risk factor for chronic traumatic encephalopathy (CTE), the production and accumulation of brain Aβ species are still considered as the main pathophysiological link between TBI and AD. Our study is the first to establish an mTBI model in 5xFAD mice and described the cognitive impairment and amyloid pathologies in the subacute phase following mTBI. We overserved that amyloid pathology and hippocampal dependent memory impairment was exacerbated within 8 days after mTBI in 3-4 months old 5xFAD mice when compared to the sham-operated group, which was largely consistent with previous studies using different transgenic models [57]. Aβ peptides are produced by neurons and other cell types, and they are subsequently eliminated via several clearance pathways including receptor-mediated transport across the BBB to the peripheral circulation [58], enzyme-mediated Aβ proteolytic degradation and removal by glial cells [50], and the perivascular drainage pathway for passive diffusion that connects to the meningeal lymphatic system [59]. Therefore, our data suggest that mTBI may alter the balance between Aβ production and clearance in the brain.

The cerebrovascular system plays a key role in removing metabolic waste products and toxic misfolded proteins such as Aβ from the brain [25]. As Aβ homeostasis sustained by vascular clearance pathways are integrated functions of normal cerebrovascular structure and an intact BBB [11,50], the loss of vascular integrity could play a key role in AD pathogenesis, as well as the mediation of tissue damage after TBI. While most current studies point to vascular injury after TBI, it remains unclear whether vascular impairment and BBB disruption induced by TBI affects the balance between amyloid production and clearance in the brain, and lead to cognitive impairment and dementia. Our study in 5xFAD mice demonstrated that mTBI exacerbates microvascular injury and BBB dysfunction in addition to existing vascular dysfunctions. Decreased BBB expression of amyloid receptors including Lipoprotein-related receptor protein 1 (LRP1) and P-glycoprotein [60] have been previously reported in TBI rodent models, suggesting increased brain amyloid production [57] and decreased transvascular clearance occur simultaneously after TBI, which is consistent with our measurement of BBB efflux showing increase brain retention of synthetic Aβ. When examining the perivascular pathway in our mTBI model, we found nearly complete blockage of the perivascular drainage within days after mTBI as well as increased retention of injected fluorescently labelled Aβ in cortical regions affected by mTBI. Nevertheless, future studies are still required to systematically determine the imbalance between Aβ production, perivascular drainage and transvascular clearance for a better understanding of TBI-induced Aβ pathogenesis and increased risk for AD, and whether vascular-directed therapies to restore BBB integrity and clearance functions can reduce the risk of TBI-related neurodegenerative diseases.

## Conclusion

In conclusion, our study demonstrated that mTBI can induce BBB dysfunction, impair the perivascular structure and the clearance of Aβ, and exacerbate amyloid pathologies and cognitive decline in mice.

## Abbreviations

AD: Alzheimer’s disease;
Aβ: β-amyloid;
ALS: amyotrophic lateral sclerosis;
ANOVA: analysis of variance;
BBB: blood-brain barrier;
CBF: cerebral blood flow;
CD13: Cluster of differentiation 13;
CNS: central nervous system;
CTE: Chronic traumatic encephalopathy;
DAPI: 4’,6-diamidino-2-phenylindole;
DG: dentate gyrus;
DPO: days post operation;
ER: endoplasmic reticulum;
GFAP: Glial fibrillary acidic protein;
ISF: interstitial fluid;
LSCI: laser speckle contrast imaging;
LRP1: Lipoprotein-related receptor protein 1;
mTBI: mild Traumatic brain injury;
MS: multiple sclerosis;
PBS: phosphate-buffered saline;
PFA: paraformaldehyde;
ROIs: regions of interest;
sTBI: severe Traumatic brain injury;
TBI: Traumatic brain injury;
TUNEL: Terminal deoxynucleotidyl transferase dUTP nick end labeling;
VEGF: vascular endothelial growth factor;
WT: wild type

## Declarations

## Acknowledgments

The authors would like to thank the RWD Life Science for assisting the Laser Speckle Contrast Imaging, Dr. Berislav Zlokovic and Dr. Yaoming Wang for guidance and technical support, and Dr. Zhonghua Dai for careful reading of the manuscript and critical comments.

## Availability of data and materials

The data that support the findings of this study are available from the corresponding author upon request.

## Authors’ contributions

YW designed and performed experiments and analyzed data and contributed to writing the manuscript. JZ, BP, SD, XX, XG, XL, SF, HW and YY performed experiments. JFC, NSM, QM, and FGP contributed to writing the manuscript. ZZ designed experiments, analyzed data and wrote the paper. All authors read and approved the final manuscript.

## Ethics approval and consent to participate

The animal experiments were approved by the Institutional Animal Care and Use Committee at the University of Southern California per NIH guidelines.

## SUPPLEMENTARY DATA

**Supplementary Fig. 1.**
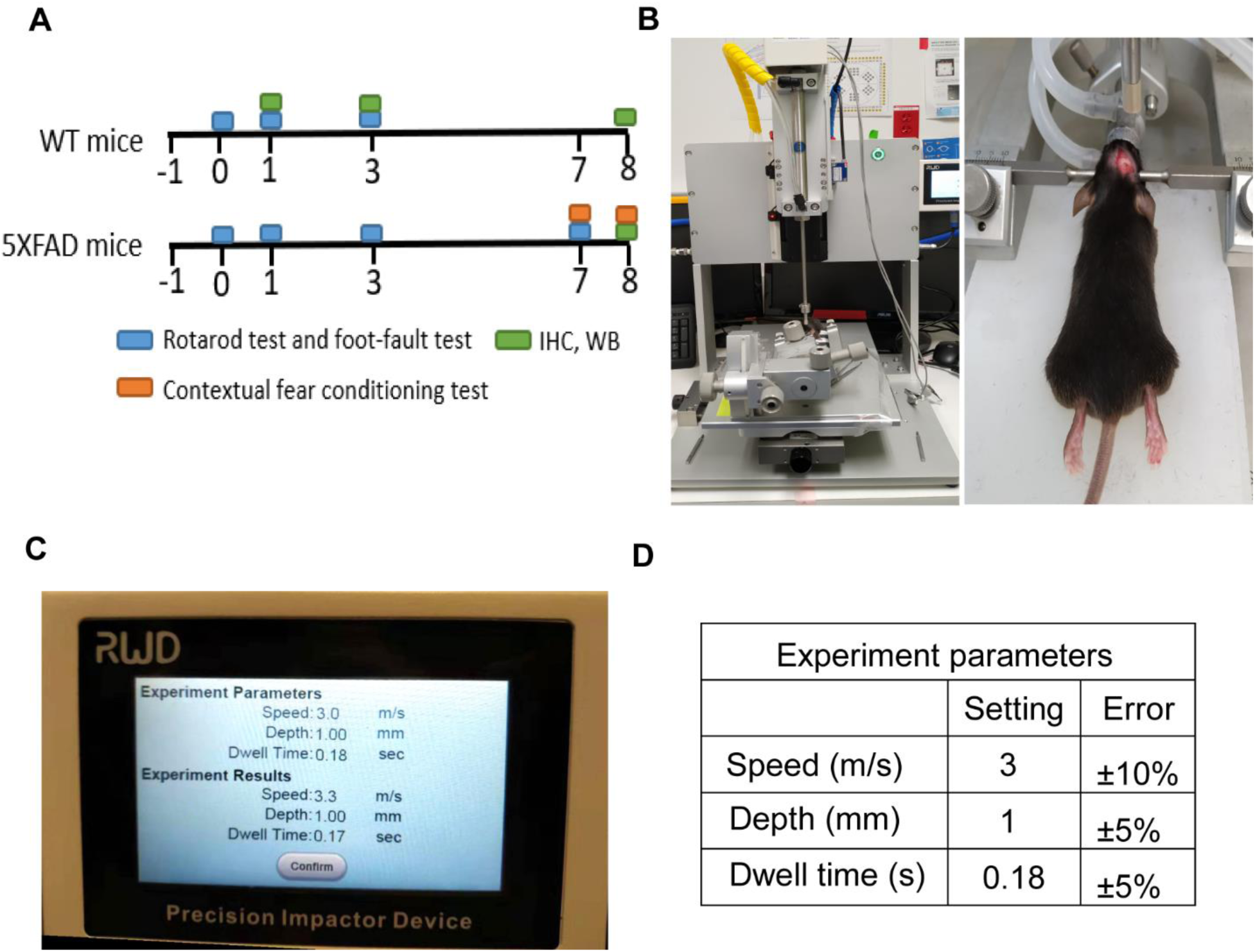
Traumatic brain injury models with controlled cortical impact in mice. (**A**) The timeline for the establishment of the TBI mouse model and the other experimental procedures. (**B**) Brain precision impactor device (RWD Life Science) to establish mTBI mouse model. (**C-D**) The experimental parameters used in the establishment of TBI mouse models.

**Supplementary Fig. 2.**
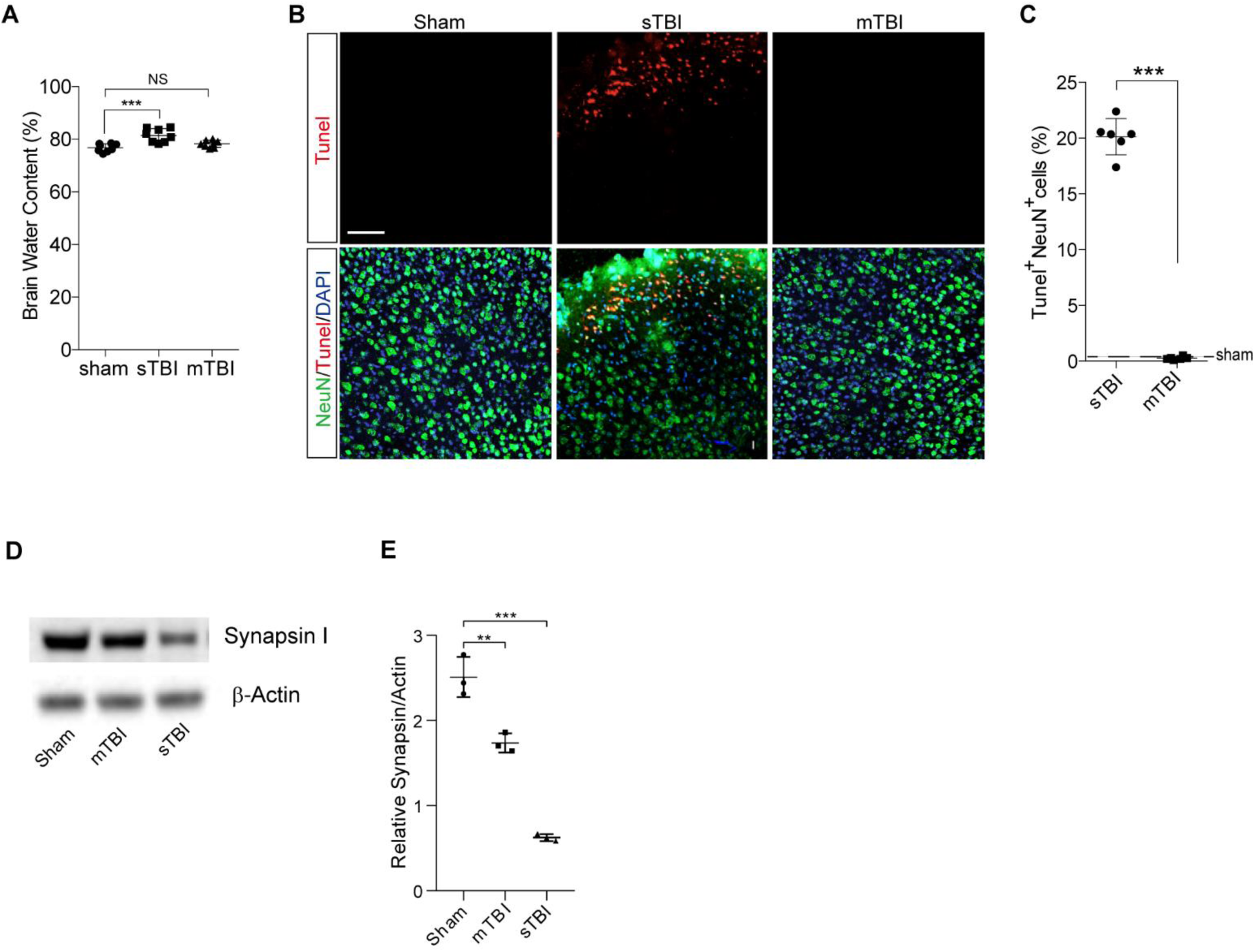
Neuropathological changes 3 days after sTBI or mTBI in mouse model. (**A**) Cerebral edema measured by brain water content. The injury significantly induced the development of cerebral edema at 24 hours post-injury in sTBI model compared to mTBI model. *n = 8* mice per group, Data are presented as Mean ± SD; ***, *P* < 0.001; NS, non-significant (*P* > 0.05), one-way ANOVA followed by Bonferroni’s post-hoc tests. (**B**) Representative confocal images showing NeuN (green) staining and TUNEL assay (red) for apoptotic neuronal death in the impacted ipsilateral area after sTBI and mTBI, or sham-operation. Scale bar, 50 μm. (**C**) Quantification of the percentage of TUNEL-positive neuronal death in sTBI (*n = 6* mice) or mTBI (*n = 6* mice). Data are presented as Mean ± SD; *** *P* < 0.001 by Student’s t-test. Dash line indicates an average value from sham-operated group (*n = 3* mice). (**D-E**) Representative immunoblots (**D**) and quantification (**E**) of Synapsin I from the cortex in the injury ipsilateral side of sham-operated, mTBI and sTBI mice. β-Actin: loading control. Data are presented as Mean ± SD; *n = 3* mice per group respectively; ***, *P* < 0.001; **, *P* < 0.01; one-way ANOVA followed by Bonferroni’s post-hoc tests.

**Supplementary Fig. 3.**
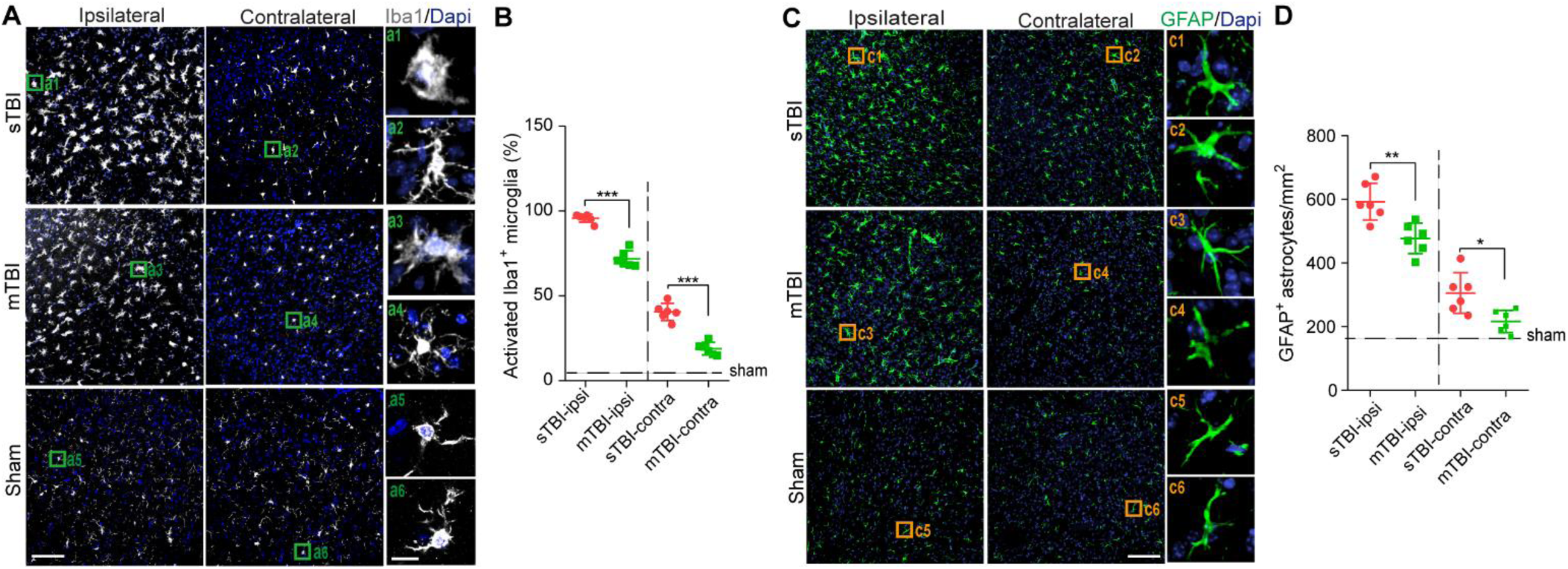
Neuroinflammatory changes 3 days after sTBI or mTBI in mouse model. (**A**) Representative images of fluorescence immunostaining for Iba1 (gray) and DAPI (blue), boxed regions show an activated and phagocytotic microglia (a1, a3) in the impacted ipsilateral side, and ramified microglia at resting state (a2, a4) in the contralateral side. Scale bar in (A) 50 μm, in (a1-a6) 10 μm. (**B**) Quantification for the percentage of activated Iba1 positive cells (*n = 6* mice per group) in the impact side (ipsi) and contralateral side (contra) in sTBI and mTBI mice (see method). Dash line indicates the average value from the sham-operated group (*n = 5* mice). (**C**) Representative images of fluorescence immunostaining for GFAP (green) and DAPI (blue), boxed regions (c1-c4) show astrocytes in the impact side and the contralateral side. Scale bar in (C) 50 μm, in (c1-c6) 10 μm. (**D**) Quantification for the GFAP positive cells per mm^2^ (*n = 6* mice per group) in the impact side (ipsi) and contralateral side (contra) in sTBI and mTBI mice. Dash line indicates the average value from sham-operated group (*n = 5* mice). Data are presented as Mean ± SD; ***, *P* < 0.001; **, *P* < 0.01; *, *P* < 0.05; NS, non-significant (*P* > 0.05) by by Student’s t-test.

**Supplementary Fig. 4.**
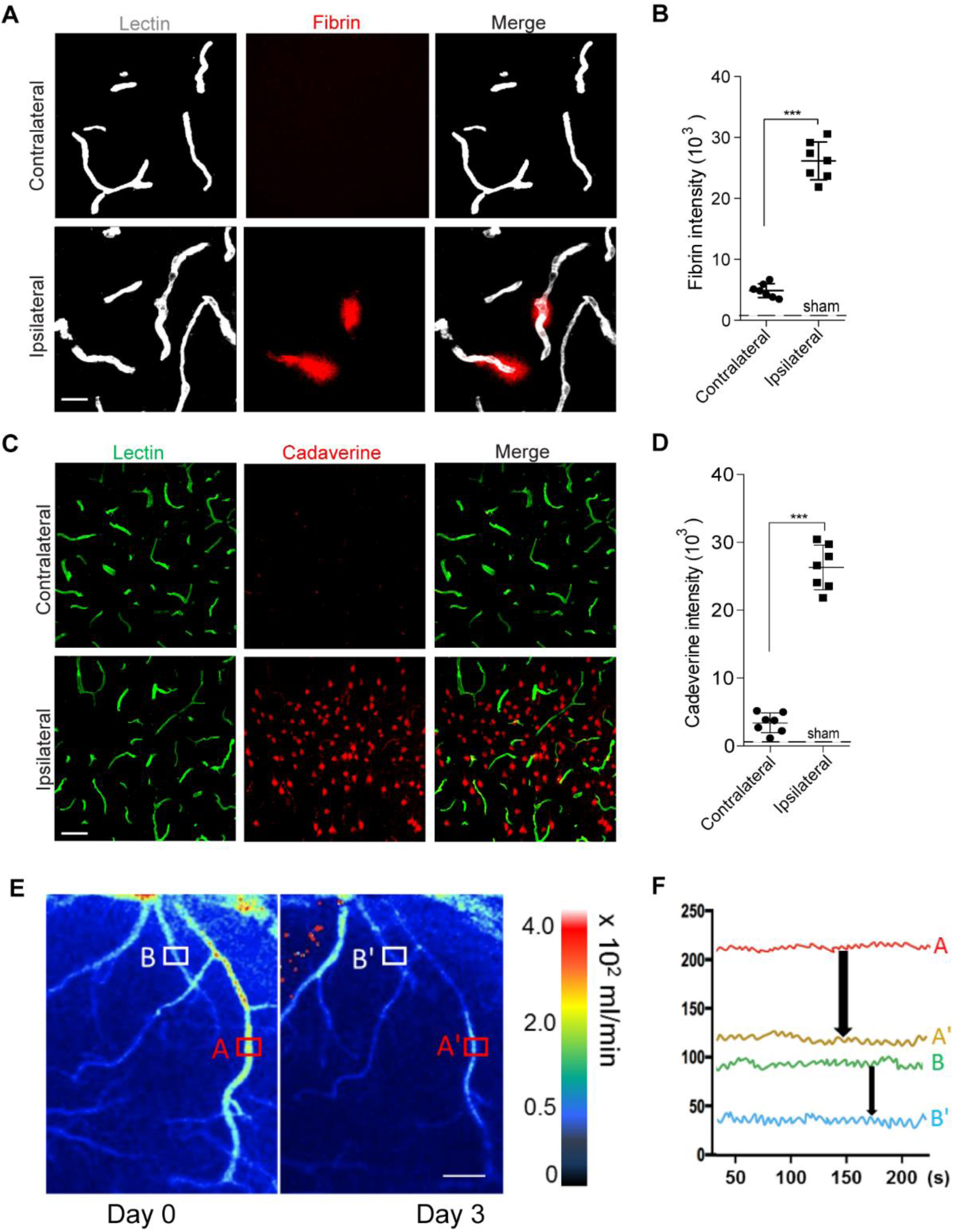
Vascular impairment in the subacute phase of mTBI in mice. (**A-B**) Representative images (**A**) and quantification (**B**) showing extravascular fibrin deposits in brain parenchyma 3 days after mTBI. Lectin (gray), fibrin (red). *n = 7* mice; Scale bar = 20 µm. (**C-D**) Representative images (**C**) and quantification (**D**) showing extravasation of intravenously administrated Alexa-555 cadaverine (red) in brain parenchyma 3 days after mTBI. Lectin (green), *n = 7* mice, scale bar = 50 µm. In B, D, data are presented as Mean ± SD; ***, P < 0.001 by Student’s t-test. Dash lines indicate average value from sham-operated group (*n = 3* mice). (**E-F**) Cerebral blood flow reduction 3 days after mTBI. Representative laser speckle contrast imaging (LSCI) images (**E**) and quantification of boxed regions (**F**) showing blood flow changes in the cortical regions 3 days after mTBI. Scale bar = 50 µm

**Supplementary Fig. 5.**
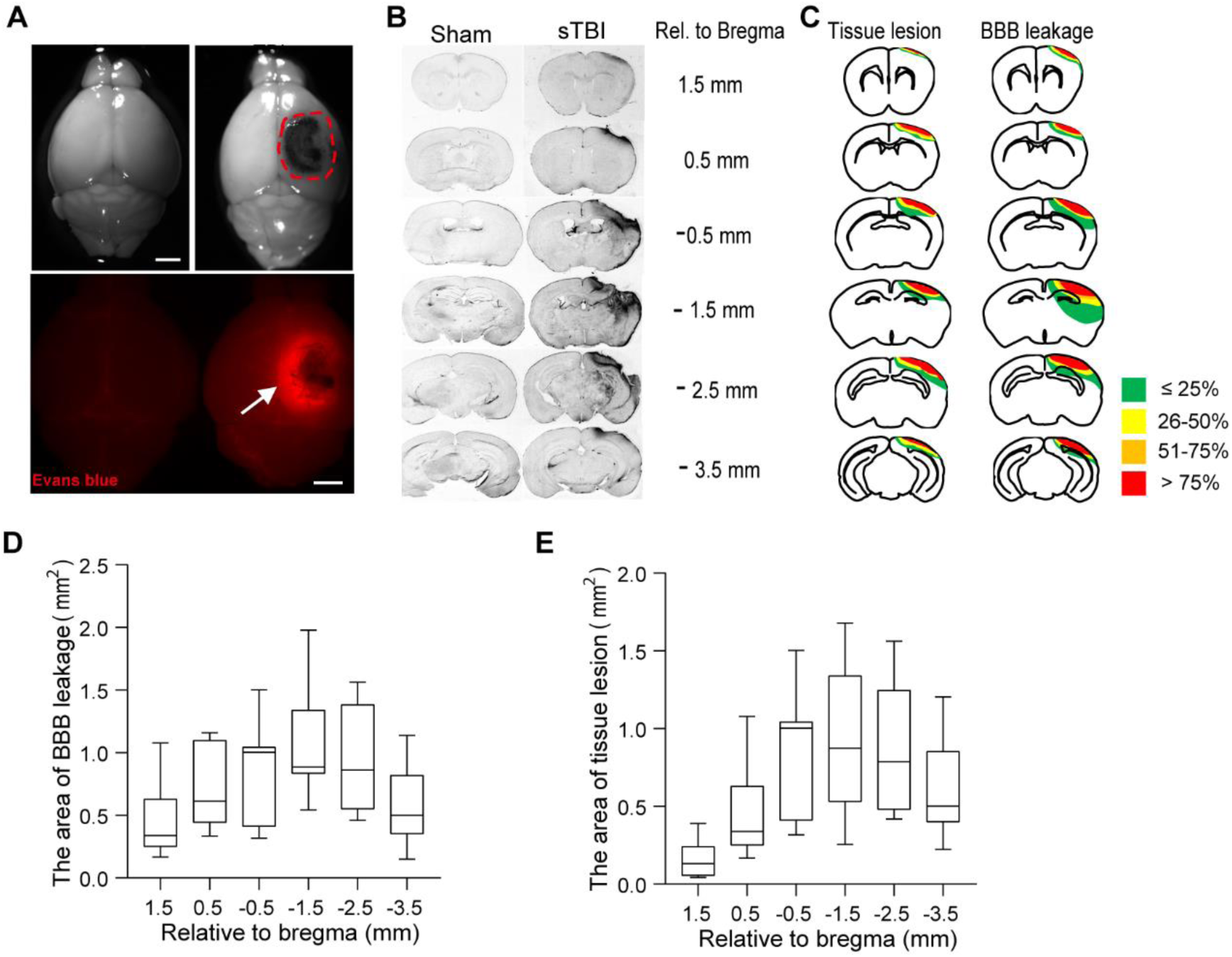
BBB dysfunction in severe TBI (sTBI) mouse model. (**A**) Evans blue extravasation and tissue loss 1-day post-injury were detected under both brightfield microscope (upper) and fluorescent microscope (below), indicating blood-brain-barrier (BBB) disruption in severe TBI mice, compared to the sham-operated mice. Red dashed circle and white arrow indicate the area of BBB leakage. Scale bar = 1 mm. (**B**) Representative coronal brain sections from (**A**) every 1 mm at 1 day after sTBI in both sham-operated mice and sTBI mice. (**C**) Accumulative map of the BBB disruption (left) and tissue lesion (right), 1 day after sTBI. Each slide depicts the accumulated information of BBB distribution and tissue lesion (color coded as indicated) at the given section in relation to the bregma. (**D-E**) The whisker plot for the area of BBB leakage based on Evans blue extravasation (**D**) and the area of tissue lesion (**E**) in brain parenchyma 1 day after sTBI, *n = 10* mice. Data are presented in box plots showing the upper (75%) and lower (25%) quartiles, with the horizontal line as the median and the whiskers as the maximum and minimum values observed

**Supplementary Fig. 6.**
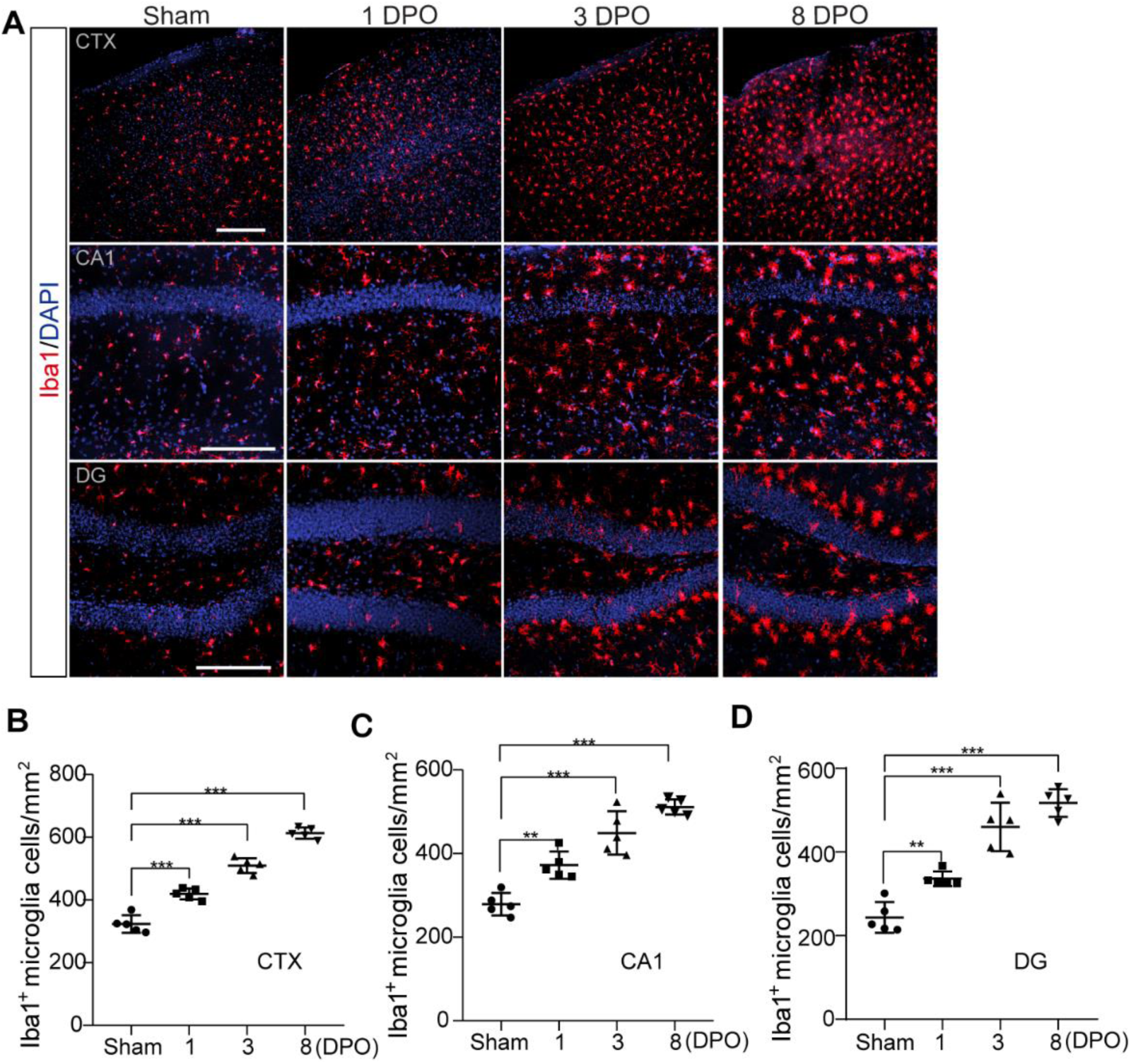
The time course of microglial activation after mTBI in mice. (**A**) Representative confocal microscope images showing Iba1-positive microglia cells (red) in cortex (CTX), Cornu Ammonis 1 (CA1) and dentate gyrus (DG) area 1-day, 3- and 8-days post operation (DPO). Scale bar = 100 µm. (**B-D**) Quantification for the Iba1-positive microglia cells per mm^2^ (*n = 5* mice per time point) in CTX, CA1 and DG area 1 day, 3- and 8-days post operation (DPO). Data are presented as Mean ± SD; ***, *P* < 0.001; **, *P* < 0.01; one-way ANOVA followed by Bonferroni’s post-hoc tests.

**Supplementary Fig. 7.**
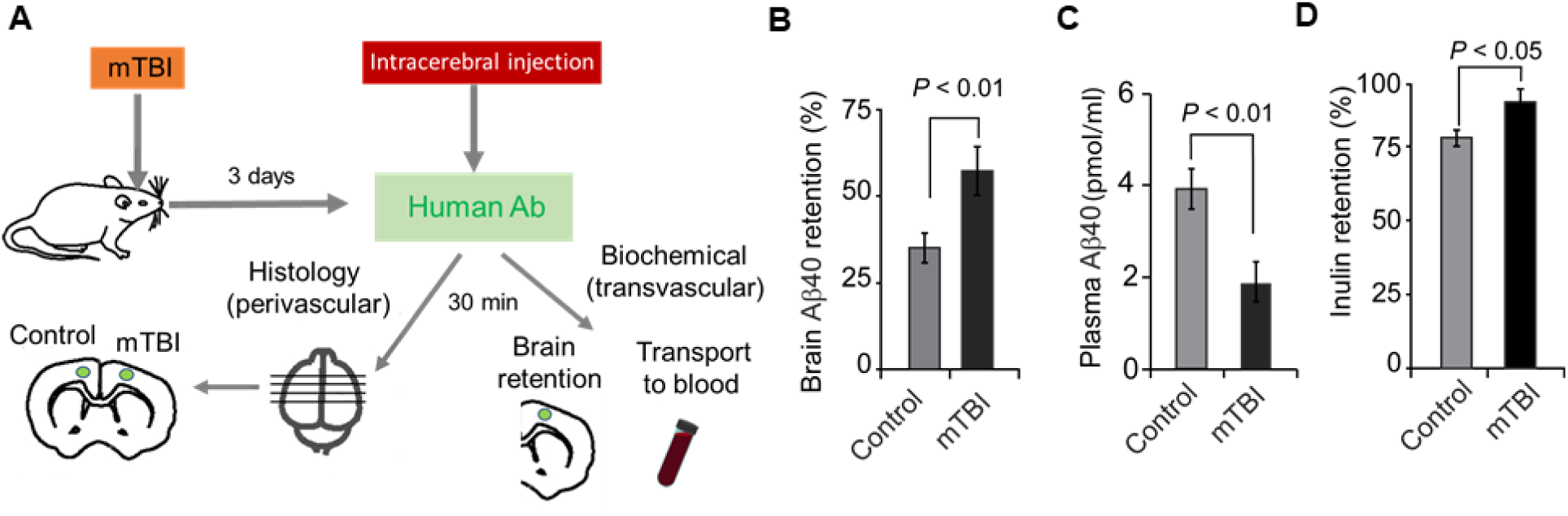
Impairment of vascular clearance pathways in the mTBI model. (**A**) Experimental procedures of *in vivo* tracing to determine transvascular and perivascular pathways for amyloid clearance in mTBI. (**B-D**) brain retention of human Aβ40 (**B**), or Aβ40 transported across the BBB into the circulation (**C**), or in brain retention of 14C-inulin (**D**), in the mTBI-affected ipsilateral hemisphere and unaffected contralateral hemisphere (control). *n = 3* mice per group; mean ± SD; *P* < 0.05; *P* < 0.01 by Student’s t-test.

